# Rumen protozoa are a hub for diverse hydrogenotrophic functions

**DOI:** 10.1101/2023.12.17.572056

**Authors:** Ido Toyber, Raghawendra Kumar, Elie Jami

## Abstract

Ciliate protozoa are an integral part of the rumen microbial community involved in a variety of metabolic processes. These processes are thought to be in part the outcome of interactions with their associated prokaryotic community. For example, methane production is facilitated by interspecies hydrogen transfer between protozoa and archaea. We hypothesize that ciliate protozoa are host to a stable prokaryotic community dictated by specific functions they carry. Here we modify the microbial community by varying the forage to concentrate ratios and show that, despite major changes in the prokaryotic community, several taxa remain stably associated with ciliate protozoa. By quantifying genes belonging to various known reduction pathways in the rumen, we find that the bacterial community associated with protozoa is enriched in genes belonging to hydrogen utilization pathways and that these genes correspond to the same taxonomic affiliations seen enriched in protozoa. Our results show that ciliate protozoa in the rumen may serve as a hub for various hydrogenotrophic functions and a better understanding of the processes driven by different protozoa may unveil potential role of ciliates in shaping rumen metabolism.

## Introduction

The rumen microbial community is considered to be one of the most diverse and complex among host associated communities ^1^. Unlike monogastric animals in which microbial fermentation represents but a fraction of the total energy balance of the host, the rumen microbiome is responsible for the vast majority of the carbon and nitrogen requirements of the ruminant animal ^2^. This is performed by a vast microbial community encompassing all domains of life; bacteria, archaea and microbial eukaryotes. The feed ingested by the host is broken down by the microbes involving complex cascades of cross-feeding and interactions that lead to the production of end products for the animal in the form of volatile fatty acids as well as the production of the unusable methane. Although the understanding of the rumen microbial ecology as well as the role and interactions between the rumen bacterial community has increasingly been studied, the eukaryotic portion of the rumen microbiome, their dynamics in the rumen and interactions remains largely neglected ^3–5^.

The largest fraction of the rumen eukaryotic community are the ciliate protozoa, which account for 2% of all microbial species and can account for up to 50% of the biomass ^6,7^. Several studies, such as Henderson et al, surveyed ruminant species across different geographies and diets, emphasized that ciliate protozoa may be more diverse than originally expected and that they may not be constrained by the same ecological forces as the prokaryotic community ^8,9^.

In addition to their still elusive ecological dynamics, ciliate protozoa have also been suggested to have many metabolic and physiological functions in the rumen, most of which not yet elucidated ^10^. Some of these functions are further suggested to be a direct consequence of their interactions with the prokaryotic community ^11,12^. The most studied functional interaction is their role in enhancing methane production via interspecies hydrogen transfer to methanogenic archaea ^12^. Such interaction was observed across a wide range of *in-vivo* and *in-vitro* studies, with the estimated enhancement of methane due to the presence of protozoa ranging from 11% to 37% ^7,12–14^, and not limited to the rumen environment ^15,16^. Additional evidence for the interaction stems from the physical association observed between ciliate protozoa and methanogens ^17–19^. These studies note that, in addition to methanogens, protozoa likely harbor a diverse bacterial community for which as of yet no type of interaction has been proposed.

In this study, we investigate the effect of host diet on the protozoa composition and the composition of their associated prokaryotic community. We find that the protozoa community is less affected by dietary changes compared to the prokaryotic community. We also discover that while the protozoa associated prokaryotic community is dependent on the free-living community and thus changes with diets, the protozoa microenvironment retains a unique community of enriched taxa independently of their abundance in the free-living community. The enrichment of these taxa may be the result of functional interactions similar to methanogens, driven by their production of hydrogen.

## Results

### Abundance and stability of the protozoa community across host animal diets

The diet of an animal encompasses a controllable determinant of changes in ruminal prokaryotic composition ^3,20–24^. However, its effect on the protozoa community and its associated prokaryotes has not been investigated. Here we varied the animal feed by changing its fiber to concentrate ratios and followed the changes occurring in the protozoa community and their associated prokaryotes (Figure 1a). To this end three groups of five cows were allotted specific feeds for a minimum of two months before sampling, with one group referred to as high fiber (HF) receiving 80/20 fiber to grain ratio, medium fiber (MF) with 50/50 ratio and low fiber (LF) receiving 30/70 ratio. After the habituation period, rumen fluid was sampled and used to characterize the abundance and composition of bacteria, archaea and protozoa. In order to characterize the protozoa-associated prokaryotic community, a subset of the fresh samples was also used to separate the protozoa community from the free-living prokaryotic community (Figure 1a).

**Figure 1.**
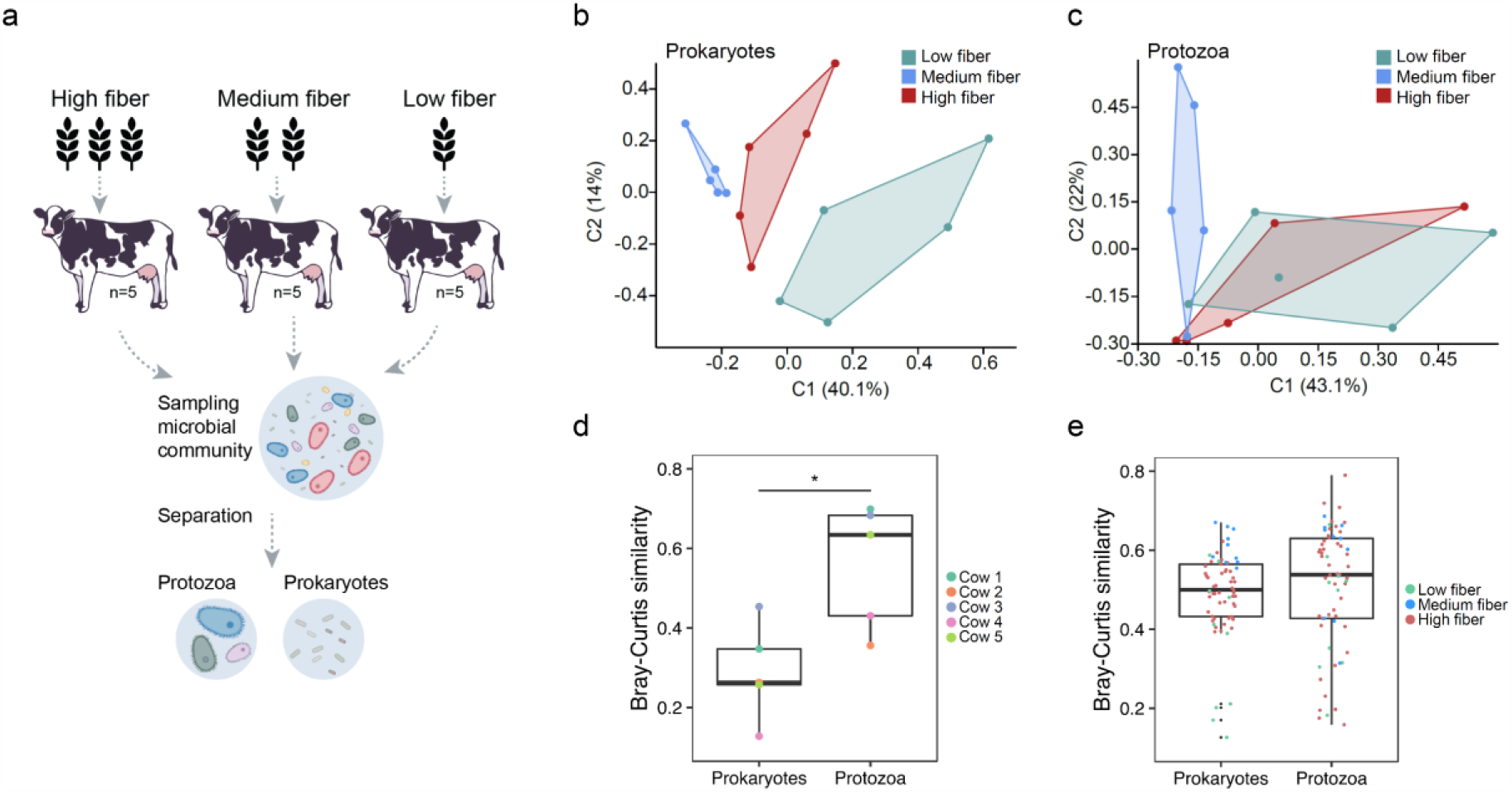
Comparison of the ecological dynamics of prokaryotes and protozoa across diets. (a) Scheme of the experimental set-up used in this study. (b), (c) Principal coordinate analysis (PcoA) based on the Bray-Curtis pairwise similarity matrix for (b) prokaryotes and (c) ciliate protozoa. Each point represents a different sample plotted according to their amplicon sequence variant (ASV) composition and abundance. A greater distance between two points infers a lower similarity in community compositions. The different colors represent the different animal diets (d) Bray-Curtis similarity between samples taken from the same five cows consecutively fed a different diet. Each point represents community similarity in each cow between the sample taken while fed high fiber and the sample taken while under low fiber diet, for the prokaryotic and protozoa community. Significance was obtained using Mann-Whitney U (*p* < 0.05). (e) Overall community similarity between samples from different cows (within diets) for the prokaryotic and protozoa community

Quantification of the abundance of ciliate protozoa across the different diets revealed that an increase in fiber content results in a decrease in the number of protozoa. This was observed when the analysis was performed using both quantitative PCR and live counting of the protozoa cells (Figure S1). The mean visual count for the HF (n = 5), MF (n = 5) and LF (n = 5) were 1.09 × 10^5^, 1.53 × 10^5^ and 2.91 × 10^5^ respectively. We did not find significant differences between archaea and bacterial abundances across the different diets (Figure S1).

We subsequently analyzed the composition of the prokaryotic and protozoa community using 16S and 18S rRNA amplicon sequencing, respectively. In line with previous observations ^20–24^, the diet of the host animal had a pronounced effect on the prokaryotic community that was observed by a significant clustering of the samples based on diet using the Bray-Curtis pairwise dissimilarity metric (ANOSIM, R = 0.709, P < 0.001, Figure 1b). In contrast, the different diets had only a subtle effect on the protozoa community (ANOSIM, R = 0.185, P = 0.054 Figure 1b), and only showed a significant difference between the HF and MF groups (ANOSIM, R = 0.21, P = 0.034).

In order to test whether the protozoa community composition is more stable compared to the prokaryotic community when direct diet change occurs, the cows under an LF diet were switched to a HF diet, and sampled again after a two months adaptation period after the diet switch occurred. We monitored the degree of changes in microbial composition occurring within each individual animal. We observed that the protozoa community varied significantly less with dietary change compared to the prokaryotic community (Student’s *t*-test, P = 0.028, Figure 1c). The mean similarity within hosts in the prokaryotic community was 0.28 compared to 0.56 for the protozoa community, the latter not significantly different compared to the natural variation between cows within a given diet (Figure 1c). It is important to note that overall, the protozoa community was not more similar between samples (within the same diet) compared to the prokaryotic community (Figure 1d). This reduces the possibility that these results stem from a naturally more similar protozoa community due to the lower species richness of protozoa..

### Stability of the protozoa associated prokaryotic community across diets

As the prokaryotic community was different between the different diets, we sought to assess whether this change in composition is also reflected in the protozoa associated community. The higher stability of the protozoa community across diets led us to hypothesize that the prokaryotic community associated with protozoa may serve as a reservoir for taxa thereby remaining present in the rumen across differing conditions. To characterize the prokaryotic community that is physically associated with protozoa, the protozoa community was separated from the free-living community in the samples taken from cows under the different diets (Figure 1a). The samples were also separated into sub-communities in order to further assess whether different protozoa taxa carry different prokaryotic communities. Our results showed that the prokaryotic community associated with protozoa (protozoa associated = PA) exhibits corresponding, significant differences in composition driven by different diets based on the Bray-Curtis dissimilarity metric (ANOSIM, R = 0.615, P =0.001, Figure 2a, Table S2). However, the communities also showed discrimination between the protozoa associated community and the free-living (FL) community (ANOSIM, R = 0.57, P =0.0001, Figure 2a, Table S2). There were no observable differences between the prokaryotic sub-communities stemming from different protozoa sub-communities (Figure S2a).

**Figure 2.**
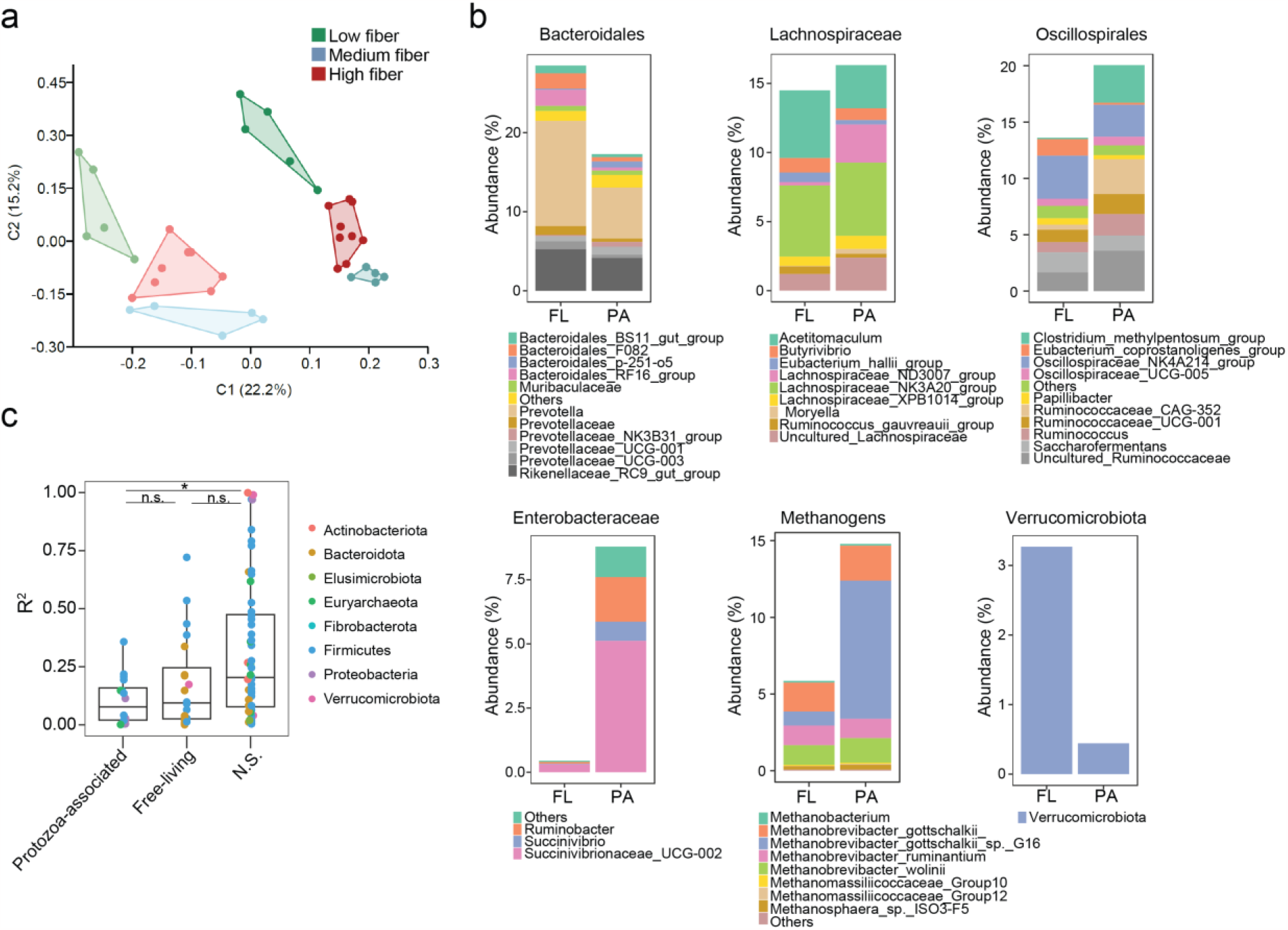
Compositional differences between the free-living and protozoa-associated prokaryotic community across diets. (a) Principal component analysis (PcoA) based on the pairwise similarity matrix obtained using the Bray-Curtis similarity index for the prokaryotic community. The different colors represent the different animal diets. The darker shades of each color represent the free-living community and lighter shades the protozoa associated community. (b) Genus level community composition between the free-living and protozoa associated prokaryotic community. (c) Box plot denoting the distribution of coefficient of variation between the free-living prokaryotic abundance and protozoa associated prokaryotic abundance.

Thus, while diet imposes an effect on the prokaryotic community structure associated with protozoa, the community composition remains unique to the protozoa association.

### Protozoa-associated community are enriched for specific taxa regardless of diet

Our results so far showed that while the prokaryotic community associated with protozoa is affected by the dietary change, it still diverges significantly from the free-living community. We therefore hypothesize that the divergence is a result of key taxa that adapted to the protozoa microenvironment.

At the phylum level, the protozoa-associated community was significantly enriched in α and γ - Proteobacteria (FL = 0.03% vs. PA = 0.42% and FL = 0.44% vs. PA = 7.9%, respectively), and Euryarchaeota (FL = 5.4% vs. PA = 14.9%), while Bacteroidetes and Verrucomicrobia were significantly more prevalent in the free-living communities (FDR corrected paired Wilcoxon, *p* < 0.01; Figure 2b, Table S3). Further characterization of the differentially abundant taxa among the protozoa-associated communities revealed that *Ruminococcaceae_CAG-352* (FL; 0.46% vs PA; 3.1%) and Clostridiales [methylpentosome] (FL; 0.09% vs PA; 3.3%) belonging to *Oscillospiraceae* (Bacillota) and Lachnospiraceae ND_3007 (FL; 0.24% vs PA; 3.9%) from Lachnospirales were significantly enriched among the protozoa-associated population (FDR corrected Paired Wilcoxon *p* < 0.01; Figure 2b, Figure S2, Table S3). From the γ-proteobacteria class, *Succinivibrionaceae, Ruminobacter* and *Succinivibrionaceae UCG-00*2 were all significantly enriched in the protozoa associated prokaryotic community. The protozoa associated communities were also shown to uniquely harbor a narrow number of ASVs belonging to the taxa mentioned such as ASVs belonging to *Methanobrevibacter, Succinovibrionaceae*, Clostridiales [methylpentosome] and Rickettsiales. Within the *Methanobrevibacter* genus, there were several ASVs detected, and associated with *M. gottschalkii sp*.*16* based on the Rumen and Intestinal Methanogen database (RIM-DB) ^25^ that were exclusive to the protozoa associated communities and those represented the majority of the methanogens found (Figure 2b, Table S3). Thus, the protozoa community harbors a subset of prokaryotic taxa that are unique or enriched regardless of the environmental conditions and the composition of the free-living community prevailing.

In order to further determine the independent nature of the enriched taxa found associated with protozoa we assessed the coefficient of determination using linear regression obtained between the abundance of the enriched taxa associated with protozoa and their relative abundance in the free-living fraction. This determines, in addition to the enrichment, whether the relative abundance found is a direct result of the change in abundance in the free-living community (between diets for instance). This analysis showed that the R^2^ of significantly enriched prokaryotic ASVs in protozoa was significantly lower than those who were not significantly associated with protozoa, suggesting that the enrichment of the specific taxa enriched in the protozoa associated community is independent of the relative abundance in the free-living community (Figure 2c). Our results thus show that a subset of taxa preferentially associates with the protozoa community and this association is stable across diets.

### The protozoa associated prokaryotic community is enriched in genes belonging to hydrogenotrophic pathways

Similar to previous studies our results reveal the association between protozoa and the methanogen community (Figure 2b) ^17,18^. This association is suggested to be the result of the release of hydrogen by the protozoa and its direct consumption by the hydrogenotrophic methanogens to produce methane ^7,12^. We thus hypothesize that hydrogen may be a strong determinant of microbial associations with protozoa and that protozoa may encompass a hub for diverse hydrogenotrophic functions beyond those observed for methanogens ^26–28^. Interestingly, we observed that taxa such as O*scillospiraceae* and *Lachnospiraceae* were significantly enriched in the protozoa-associated prokaryotic community, which are taxa that have been previously shown to utilize hydrogen for reductive acetogenesis and the Wood Ljungdahl pathway ^29–31^.

In order to test whether hydrogenotrophic functions are enriched associated with protozoa, we investigated the presence of other bacterial reductive pathways known to be prevalent in the rumen. To do this we quantified key genes of pathways related to hydrogen utilization and reduction of end molecules known to be present in the rumen and evaluated their abundance in the protozoa associated community compared to the free-living community.These genes included the formyltetrahydrofolate synthetase genes and acetyl-COA synthase (*ftfhs, acsB*) of the Wood Ljungdahl pathway ^30^, the adenosine-5′-phosphosulfate reductase (*aprA*) and the dissimilatory sulfite reductase subunit alpha (*dsrA*) ^32,33^ of the dissimilatory sulfate reduction pathway, the methyl coenzyme-M reductase (*mcrA*) ^34^, and the nitrite reductase (nrfA), of the dissimilatory nitrate reduction pathway. Our results showed that the *mcrA* gene was the most abundant gene in our samples and was significantly higher in the protozoa associated communities compared to the free-living community (Figure 3a, b), which is in line with the higher proportion of methanogens reads observed using the amplicon sequencing (Figures 2b). The *ftfhs* and *acsB* gene, belonging to the Wood Ljungdahl pathway were generally the second most abundant genes observed but were not significantly enriched in the protozoa associated prokaryotic community compared to the free-living community (Figure 3a). The *nrfA* gene of the dissimilatory nitrate reduction pathway was significantly enriched in protozoa associated bacterial communities reaching up to ten-fold higher abundance compared to the free-living community (paired Wilcoxon; *p* <0.01, Figure 3a). Such enrichment was also seen in the genes *aprA* and *dsrA* belonging to the dissimilatory sulfate reduction pathway (paired Wilcoxon; *p* <0.05, Figure 3a). Interestingly, the diet had an effect on gene abundance, with the most consistent feature being an increase in the gene *nrfA* proportion concomitant with an increase in fiber content in the diet in the protozoa associated community (Figure 3b). Our results overall show that specific bacterial hydrogenotrophic activity may be higher among the protozoa-associated prokaryotic community compared to the free-living community supporting hydrogen as an important driver of microbial interactions between protozoa and prokaryotes.

**Figure 3.**
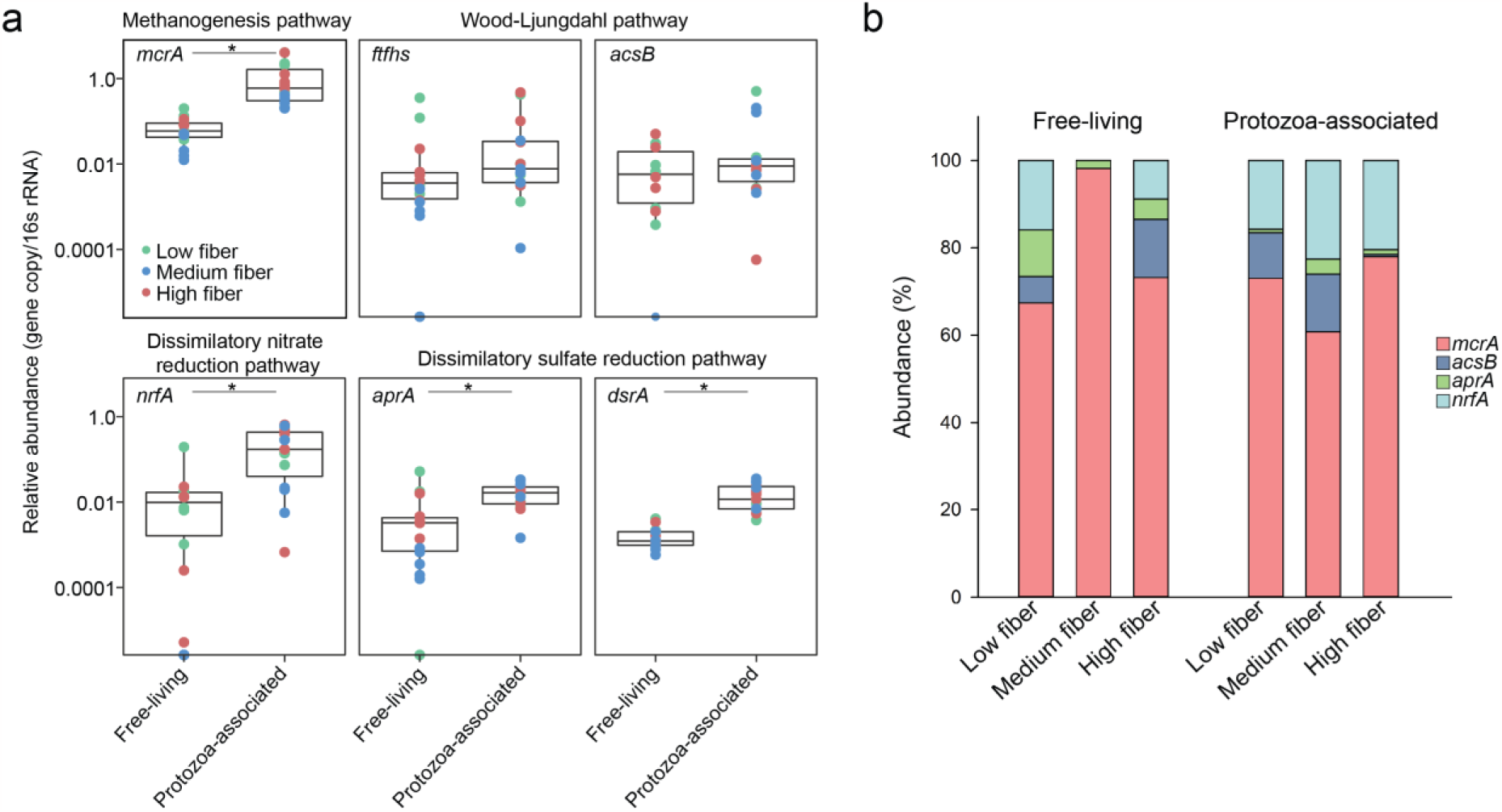
Relative abundance of key genes in hydrogenotrophic pathways in the free-living and protozoa associated prokaryotic community. (a) Box plots illustrating the abundance of genes belonging to known hydrogenotrophic pathways in the rumen in both free-living and protozoa-associated prokaryotic communities of the rumen. Significance between the groups was assessed using a paired Wilcoxon statistical test with false discovery rate (FDR) correction (p < 0.05). (b) Proportion of genes representing these pathways observed across various diets within the free-living and protozoa-associated prokaryotic communities of the rumen.

### Phylogenetic assignment of the hydrogen utilizing functions

Our results show the presence and stability of specific prokaryotic taxa with the protozoa environment (Figure 2) as well as an enrichment in specific hydrogenotrophic functions (Figure 3). In order to integrate the specific taxa with the enrichment in functions, the amplified gene sequences from the different hydrogenotrophic pathways were cloned and sequenced. The gene sequences were then compared to curated metagenome assembled genomes (MAGs) datasets obtained from large scale surveys of the rumen microbiome ^35,36^, alongside previous sequencing efforts obtained for the different hydrogenotrophic functions in the rumen and bacterial isolates known to carry the pathways ^29,30^. Out of eleven sequences obtained from the *ftfhs* gene, ten had a closest match to Clostridiales MAGs, and more specifically, to Lachnospiraceae, with sequence similarities ranging between 69-99% to their closest relative (Figure 4a, Table S4). This result also matches the enrichment of these taxa observed through the 16S rRNA analysis (Figure 2b). We further assessed whether the full Wood-Ljungdahl pathway was present in the closest related MAGs to our sequences (Figure 4). For the case of the *ftfhs* gene, only one of the sequences obtained from the protozoa associated gene amplification was similar to a MAG carrying both the *ftfhs* and the *acsB* genes (Figures 4a). The *acsB* sequences in contrast were associated with MAGs carrying both genes and had generally a more complete Wood-Ljungdahl pathway. Notably, only one Lachnospiraceae MAG associated with many sequences from *ftfhs* and *acsB* genes could be found. This strengthens previous observations that *acsB* may constitute a better marker for the assessment of reductive acetogenesis ^29,37^.

**Figure 4.**
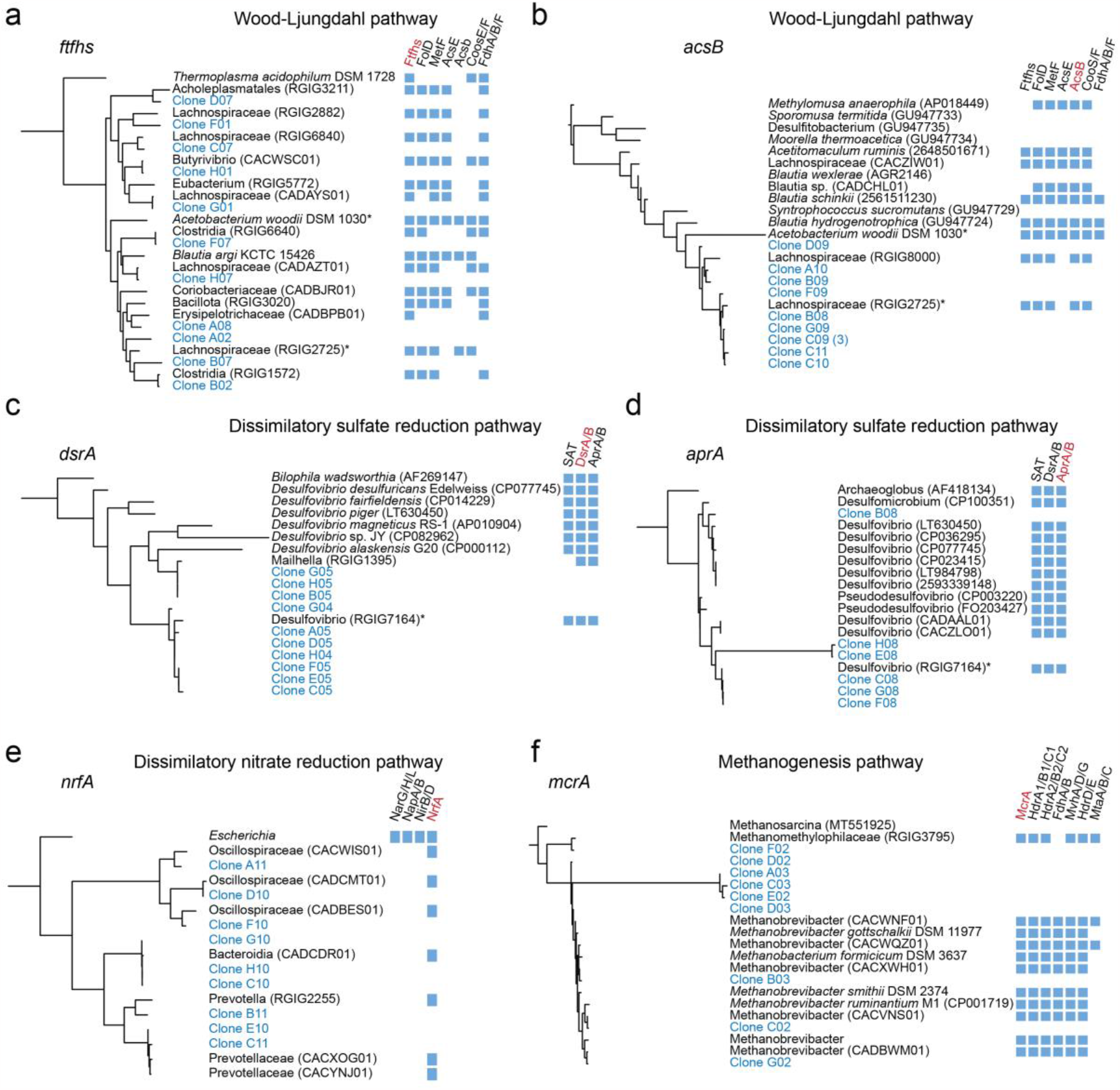
Phylogenetic assignment of the gene sequences and complementarity of hydrogenotrophic pathways. Phylogenetic trees of gene sequences obtained in this study (in blue), MAGs obtained from previous large scale metagenomic analysis (accessions starts with “CA” or “RGIG” 35,36), and sequences form the NCBI database. The genomes of homologs were searched against genes of the Wood-Ljungdahl pathway (a, b), the dissimilatory sulfate reduction pathway (c, d), the dissimilatory nitrate reduction pathway (e) and the hydrogenotrophic methanogenesis pathway (f). The presence of specific genes is indicated by blue squares. Species or MAGs that are shared between genes from the same pathway are marked with an asterisk.

All the *dsrA* and *aprA* genes from the dissimilatory sulfate reduction pathway were attributed to MAGs of the Desufovibrionaceae family, with a shared Desulfovibrio MAG (Figure 4c, d). Interestingly, although not identified by the original statistical analysis of the 16S rRNA data (which considered only ASVs and genera exhibiting a relative abundance higher than 0.5%), this family was in a significantly higher relative abundance in association with protozoa (Figure S3). For the ammonia-forming nitrite reductases (*nrfA*), the gene sequences matched MAGs belonging to *Prevotellaceae* (6 clones) and *Oscillospiraceae* (4 clones) and was the only gene present from the dissimilatory nitrate reduction pathway in the closely related MAG sequences.

## Discussion

The ecological dynamics of ciliate protozoa in the rumen remains seldom analyzed when compared to bacteria and archaea. This phenomenon is not only true for the rumen environment but has been true for most other environments leaving a gap in our understanding of complex microbial communities ^4,38,39^. Our results show that while the rumen prokaryotic community responds strongly to a change in diet as a community, the protozoa do not respond to changes in their environment caused by diet. More specifically, while the protozoa population size increases with decreased fiber content in the feed as previously observed ^40,41^, there were no significant changes in community composition, as opposed to the bacterial community. Henderson *et al*, which assessed the protozoa community across different cohorts of different species of ruminants, similarly reported a strong host individuality in protozoa communities even within specific cohorts ^8^. Furthermore, a recent study in the yak rumen found several differences in the relative abundance of some protozoa species in response to feed type, but overall community structure as measured by Beta diversity was not significantly different as opposed to bacterial or the fungal community ^9^.

Our results further support the notion of a higher stability of protozoa in the rumen, which was observed by examining the community changes when the animal diet was switched (Figure 1c). Overall, we reveal a large difference in the response between protozoa and their prokaryotic neighbors to stark changes in the rumen environment such as animal diet. This observation hints to diverse unexplored ecological forces driving the community of protozoa in the rumen. The individuality in host animals may be driven by genetics factors ^42^, or alternatively, differential early acquisition may dictate protozoa composition later in life. The latter alternative may carry important implications as to the possibility of early intervention in order to stably modulate the protozoa community ^3^.

Studies on association between protists and prokaryotes in general and specifically ciliate protozoa have shown that the nature of interaction can be diverse, ranging from antagonistic interaction for one or both partners to commensal, all the way to mutualistic interactions ^38,43–45^. Our results indicate that the ciliate protozoa associated prokaryotic community is strongly driven by the community surrounding them, and changes induced in this community results in a different community. This was seen in two levels, with different diets modulating the protozoa associated bacterial community and on the individual level, as the composition reflected the free-living community in each individual host. However, our results also show that the protozoa associated community retains significant differences when compared to the free-living community regardless of diet, suggesting that a certain degree of selection may occur in this community. This is particularly evident for taxa belonging to the Proteobacteria phylum and Oscillospirales.

Protozoa are well-established hydrogen-producers, creating an attractive environment for taxa of the group of methanogens that utilizes hydrogen to reduce CO_2_ to methane ^7,12^. By quantifying key genes belonging to hydrogen utilizing pathways known to exist in the rumen, we found that the community associated with ciliate protozoa is significantly enriched in those genes. In the case of potential nitrite and sulfate reducing taxa, the gene sequences obtained could be attributed to taxonomic affiliations seen as enriched in association with protozoa via amplicon sequencing. Our results thus suggest that ciliate protozoa in the rumen may be a hub for various hydrogenotrophic functions, beyond methanogenesis. This observation is not without precedents and different types of hydrogen transfer based putative mutualism have been suggested between prokaryotes and ciliate ^43,46,47^. In termites, putative bacterial symbionts from the Myxococcota (formerly Deltaproteobacteria) and Spirochaetota phylum were shown to have a wide range of hydrogenotrophic function which appear to be the nature of the interaction between them and their flagellate protozoa host ^48–50^. Likewise, in anoxic fresh water lake, a ciliate endosymbiont was found to supply its ciliate host with energy via denitrification performed by hydrogen transfer _51_.

In the rumen, metabolic interactions between ciliate protozoa and nitrate/nitrite reducing bacteria have been proposed. When co-cultured with bacteria, rumen protozoa were reported to accelerate nitrate reduction ^11^, and the protozoal fraction, which likely included both protozoa associated prokaryotes showed a greater ability to reduce nitrate without accumulation of nitrite. The possibility that, combined nitrate reduction to nitrite by protozoa, for which genomic evidence exists, with production of hydrogen may lead to the enrichment of nitrite reducing bacteria ^52,53^. Considering the observation that exogenous addition of nitrite (but not nitrate) severely inhibits motility and chemotaxis of protozoa ^53^, we can speculate that its clearance by bacteria might encompass the mechanism of mutualistic interaction between them. Our results suggest that the protozoa associated microbiome may possibly complete the reduction of nitrate. Successful decoupling of protozoa and associated bacteria would be required to assess such. As nitrate and other nitrogenous compounds are being evaluated for their potency to reduce methane, the role of protozoa and their associated microbes in metabolizing these compounds and their contribution to the mitigation of methane reduction is an interesting avenue for further research.

Overall, our results reveal the potentially impactful role of ciliate rumen protozoa as stable interactions partners and as a source of energy and habitat for diverse prokaryotic species.

## Material and methods

### Animals handling and sampling

The experimental procedures used in this study were approved by the Faculty Animal Policy and Welfare Committee of the Agricultural Research Organization Volcani Research Center approval no. 889/20 IL, in accordance with the guidelines of the Israel Council for Animal Care.

For the experiment, 15 cows were divided into three groups by their fiber/grain ratio: Low-Fiber (LF) refers to a total mixed ration (TMR) composed of ∼70:30 grain to fiber ratio, Medium-Fiber (MF) refers a feed ration composed of a grain to fiber ratio 50:50, and High-Fiber (HF) refers to the cows receiving a diet composed of 20:80 grain to fiber ratio. Rumen fluid was collected after two months of adaptation to the feed via a custom-made stainless-steel stomach tube, connected to an electric-powered vacuum pump (Gast, Inc. MI, USA). The rumen fluid passed through eight layers of cheesecloth into CO_2_ filled glass bottles and immediately transferred to an oxygen-free environment in an anaerobic glove box (Coy Inc. MI, USA).

### Rumen sample processing

A subset of the samples was immediately frozen for overall compositional analysis of the microbiome. For the remainder of the samples meant for the assessment of the protozoa associated prokaryotic community, we used a sedimentation technique as the protozoa cells are significantly larger than prokaryotic cells, allowing an efficient separation with the free-living prokaryotic community as previously performed ^18,54,55^. Briefly, under an anaerobic environment, we transfer the rumen fluid mixed at 1:1 ratio with pre-warmed, anaerobic Coleman salt buffer to a separating funnel and incubate for 50 minutes at 39°C ^54^. Afterward, glucose was added in 1g/L concentration to increase fermentative activity in the purpose of flocculent debris to the surface and improve the separation of the protozoal sediment for additional 20 minutes ^54^. To isolate the protozoa fractions from external prokaryotic cells stemming from the free-living population, the protozoa community was washed four times by centrifugation at 500 × g for 5 min and subsequent fresh buffer replacement. The sediment from the last wash was suspended in a 3 ml extraction buffer and transferred immediately to -20°C for downstream processing and analysis.

### Protozoa visual quantification

To determine the number of protozoa in the whole rumen samples we took 1 ml of whole rumen fluid and centrifuged it at 500 × g for 5 min. Next, we suspended the protozoa pellet with a 1:10 ratio of 4% PFA at 7.4 PH and added two drops of brilliant green dye for protozoa staining and incubated for 1 hour for better staining results. The fixed 1 ml protozoa suspension was transferred to a Sedgewick Rafter counting chamber (Marienfeld, Germany), and counted the number of protozoa on each grid under an x10 microscope objective lens (Zeiss, Germany). Overall for each sample, at least 25 squares were counted and the number of protozoa was extrapolated to the number of protozoa in one ml of rumen.

### DNA extraction

DNA extraction was performed as previously described ^56^. In brief, cells were lysed by bead disruption using a Biospec Mini-Beadbeater-16 (Biospec, Bartlesville, OK, USA) at 3,000 RPM for 3 min with phenol followed by phenol/chloroform DNA extraction. The final supernatant was precipitated with 0.6 volume of isopropanol and resuspended overnight in 50-100 μl TE buffer (10 mM Tris-HCl, 1 mM EDTA), then stored at 4 °C for short-term use, or archived at −80 °C.

### Illumina amplicon sequencing

Amplicon sequencing was performed by the Hylab Research Laboratory (Rehovot, Israel). Briefly, 20 ng of DNA was used in a 25 μl PCR reaction with primers, using PrimeStar Max DNA Polymerase (Takara) for 20 cycles. The PCR product was purified using AmpureXP beads, and a second PCR to add the adapter and index sequences was performed using the Fluidigm Access Array primers for Illumina. The PCR reaction was purified using AmpureXP beads and the concentrations were measured by Qubit. The samples were pooled, run on a DNA D1000 screentape (Agilent) to check for correct size and for the absence of primer-dimers product. The pool was then sequenced on the Illumina MiSeq, using the MiSeq V2-500 cycles sequencing kit. Sequencing was performed on a MiSeq platform using the paired end protocol (2 × 250 bp). The primers sequences used for the 16S rDNA bacteria and archaea were taken from the updated sequences of the earth microbiome project, amplifying the V4 region 515F (5′-GTGYCAGCMGCCGCGGTAA-3′) and 806R (5′-GGACTACNVGGGTWTCTAAT-3′) ^57^. For the 18S rDNA sequencing, primers specifically designed for ciliates were used from Tapio et al. (2016) with the following sequences: CiliF (5′-CGATGGTAGTGTATTGGAC-3′) and CiliR (5′-GGAGCTGGAATTACCGC-3′) ^58^.

### Quantitative real-time PCR for microbial domains and key genes in hydrogenotrophic pathways

To determine the copy number of the different ruminal microbial populations and the bacterial hydrogenotrophic genes, we use the standard curve method ^59^. The curve for each sequence was obtained by first amplifying sequences of our targeted genes; the PCR products were run and extracted from a 1.5% agarose gel. Each amplification product was quantified for the number of copies and serial 10-fold dilution was made. Real-time PCR was performed in a 10 µl reaction mixture containing 5 µl of Absolute Blue SYBR Green Master Mix (Thermo Scientific, Waltham, MA, United States), 0.5 µl of each primer (10µM working concentration), 2µl of nuclease-free water and 2µl of 10ng/µl DNA templates.

The primers used throughout this experiment and the conditions for the reactions can be found in Table S1.

### Clone library sequence alignment, phylogenetic tree, and taxonomic annotation

The PCR products of the representative genes of the hydrogenotrophic pathways were run on 1.5% agarose gel and purified using Gel DNA Recovery Kit (Hylabs, Rehovot, Israel), cloned in the PGEM-T easy vector (Promega, U.S) and transformed into competent *E*.*coli* cells (DH5α). At least 10 clones per gene were picked for Sanger sequencing in order to assess the phylogenetic and taxonomic affiliation of the genes. For this analysis, we used previously performed large scale metagenomic studies with thousands of metagenome-assembled genomes (MAGs), and compared the extracted sequences to the ones obtained in this study ^35,36^. We additionally used Blast using the nr database on our sequences to uncover sequences not covered by the mentioned databases. We additionally added a comparison with sequences stemming from bacterial isolates known to carry a given pathway. All the sequences were trimmed based on the primer coverage and aligned using MAFFT (version 7) ^60^. CD-hit was used to remove redundancies in sequences at 97% similarity (except for the sequences obtained from this study) ^61^. The resulting alignment was used for the construction of a maximum-likelihood phylogenetic tree using IQTree ^62^, with LG model and 1000 bootstrap replicates. The phylogenetic tree was visualized using iTOL ^63^. We further assessed the complementarity of the pathways for each existing genome and MAGs. We additionally added to the comparison sequences stemming from bacterial isolates known to carry a given pathway.

We further assessed the complementarity of the pathways for each existing genome, and MAG used the Prokka annotation of the genome ^64^. Afterward, each genome protein annotated file was uploaded in the web-based tool KofamKOALA - KEGG Orthology Search and each KEGG hit was mapped in the KEGG map with the default threshold value and searched for pathway completeness ^65,66^.

### Data analysis

Downstream processing of the 16S rRNA data, up to the generation of the amplicon sequence variant table (ASV) was performed in QIIME v.2 ^67^. DADA2 was applied to model and correct Illumina-sequencing amplicon errors and clustering of ASVs ^68^. Taxonomic assignment for the bacterial 16S rRNA was performed using the pre-trained classifier Silva database (Silva_138.1_ssu) ASVs from 515F/806R region from QIIME v.2 pipeline ^69^. For a more accurate and rumen specific taxonomic assignment of the archaeal sequences, the Rumen and Intestinal Methanogen database (RIM-DB), which enables a deeper classification or rumen methanogenic taxa, was used to categorize the methanogenic ASVs according to defined clades ^25^.

After the generation of the ASV table, singletons/doubletons were removed and subsampling to an even depth of 4000 reads per sample was performed for all subsequent analyses. Alpha and Beta diversity analyses were performed and plotted using the PAleontological STatistics software ^70^, including principal coordinate analysis (PCOA) using the Bray–Curtis dissimilarity metric and ASV richness, evenness, and Shannon index. Analysis of similarity (ANOSIM) was used to test the significance of the group clustering.

Centered log-ratio transformation was performed for the statistical analysis of compositional data, using the ‘compositions’ R package ^71^. For most of the statistical analyses pertaining to the comparison between free-living and protozoa-associated prokaryotic we used paired Wilcoxon tests unless otherwise stated. For all the analyses, *p* values of <0.05 after false discovery rate correction via the Benjamini-Hochberg procedure were considered significant, unless otherwise stated in the text or figure. Statistical tests and data analysis across the different fractions were performed in R version 3.5.3 ^72^.

## Acknowledgements

We want to thank the ARO farmer and veterinarians for their support throughout all experiments. We thank Dr. Tanita Wein for her critical reading of the manuscript and her insights. This study was supported by grants from the Israeli Dairy Board foundation (Grant No. 362-0699/80) and the Israeli Science Foundation (Grant No. 603/20).

## Notes

### Competing Interest Statement

The authors have declared no competing interest.

## References

1. de Jonge, N., Carlsen, B., Christensen, M. H., Pertoldi, C. & Nielsen, J. L. The gut Microbiome of 54 mammalian species. Front. Microbiol. 13, 886252 (2022).

2. Bergman, E. N. Energy contributions of volatile fatty acids from the gastrointestinal tract in various species. Physiol. Rev. 70, 567–590 (1990).

3. Mizrahi, I. & Jami, E. A method to the madness: Disentangling the individual forces that shape the rumen microbiome. EMBO Rep. 22, e52269 (2021).

4. Solomon, R. & Jami, E. Rumen protozoa: from background actors to featured role in microbiome research. Environmental Microbiology Reports vol. 13 45–49 Preprint at 10.1111/1758-2229.12902 (2021).

5. Firkins, J. L., Yu, Z., Park, T. & Plank, J. E. Extending Burk Dehority’s perspectives on the role of ciliate protozoa in the rumen. Frontiers in Microbiology vol. 11 Preprint at 10.3389/fmicb.2020.00123 (2020).

6. Cottle, D. J., Nolan, J. V. & Leng, R. A. Turnover of protozoa and bacteria in the rumen of sheep. Proc. Aust. Soc. Anim. Prod (1978).

7. Newbold, C. J., de la Fuente, G., Belanche, A., Ramos-Morales, E. & McEwan, N. R. The role of ciliate protozoa in the rumen. Front. Microbiol. 6, (2015).

8. Henderson, G. et al. Rumen microbial community composition varies with diet and host, but a core microbiome is found across a wide geographical range. Sci. Rep. 5, (2015).

9. Cui, X. et al. Dynamic changes in the yak rumen eukaryotic community and metabolome characteristics in response to feed type. Front Vet Sci 9, 1027967 (2022).

10. Li, Z. et al. Genomic insights into the phylogeny and biomass-degrading enzymes of rumen ciliates. ISME J. 16, 2775–2787 (2022).

11. Villar, M. L. et al. Dietary nitrate and presence of protozoa increase nitrate and nitrite reduction in the rumen of sheep. J. Anim. Physiol. Anim. Nutr. 104, 1242–1255 (2020).

12. Ushida, K., Newbold, C. J. & Jouany, J.-P. Interspecies hydrogen transfer between the rumen ciliate Polyplastron multivesiculatum and Methanosarcina barkeri. J. Gen. Appl. Microbiol. 43, 129–131 (1997).

13. Solomon, R. et al. Protozoa populations are ecosystem engineers that shape prokaryotic community structure and function of the rumen microbial ecosystem. ISME J. (2021) doi:10.1038/s41396-021-01170-y.

14. Ranilla, M. J., Jouany, J.-P. & Morgavi, D. P. Methane production and substrate degradation by rumen microbial communities containing single protozoal species in vitro. Lett. Appl. Microbiol. 45, 675–680 (2007).

15. Treitli, S. C. et al. Hydrogenotrophic methanogenesis is the key process in the obligately syntrophic consortium of the anaerobic ameba Pelomyxa schiedti. ISME J. 17, 1884–1894 (2023).

16. Yamada, K., Kamagata, Y. & Nakamura, K. The effect of endosymbiotic methanogens on the growth and metabolic profile of the anaerobic free-living ciliate Trimyema compressum. FEMS Microbiol. Lett. 149, 129–132 (1997).

17. Lloyd, D. et al. Intracellular prokaryotes in rumen ciliate protozoa: Detection by confocal laser scanning microscopy after in situ hybridization with fluorescent 16S rRNA probes. Eur. J. Protistol. 32, 523–531 (1996).

18. Levy, B. & Jami, E. Exploring the prokaryotic community associated within the rumen ciliate protozoa population. Front. Microbiol. 9, 2526 (2018).

19. Park, T. & Yu, Z. Do ruminal ciliates select their preys and prokaryotic symbionts? Front. Microbiol. 9, 1710 (2018).

20. Shaani, Y., Zehavi, T., Eyal, S., Miron, J. & Mizrahi, I. Microbiome niche modification drives diurnal rumen community assembly, overpowering individual variability and diet effects. ISME J. 12, 2446–2457 (2018).

21. Jami, E. et al. Effects of including NaOH-treated corn straw as a substitute for wheat hay in the ration of lactating cows on performance, digestibility, and rumen microbial profile. J. Dairy Sci. 97, 1623–1633 (2014).

22. Friedman, N. et al. Diet-induced changes of redox potential underlie compositional shifts in the rumen archaeal community. Environ. Microbiol. (2016).

23. Mackie, R. I. & McSweeney, C. Improving rumen function. (Burleigh Dodds Science Publishing, 2020).

24. Newbold, C. J. & Ramos-Morales, E. Review: Ruminal microbiome and microbial metabolome: effects of diet and ruminant host. Animal 14, s78–s86 (2020).

25. Seedorf, H., Kittelmann, S., Henderson, G. & Janssen, P. H. RIM-DB: a taxonomic framework for community structure analysis of methanogenic archaea from the rumen and other intestinal environments. PeerJ 2, e494 (2014).

26. Greening, C. et al. Diverse hydrogen production and consumption pathways influence methane production in ruminants. ISME J. 13, 2617–2632 (2019).

27. Morgavi, D. P., Forano, E., Martin, C. & Newbold, C. J. Microbial ecosystem and methanogenesis in ruminants. Animal 4, 1024 (2010).

28. Li, Q. S. et al. Dietary selection of metabolically distinct microorganisms drives hydrogen metabolism in ruminants. ISME J. 16, 2535–2546 (2022).

29. Gagen, E. J., Padmanabha, J., Denman, S. E. & McSweeney, C. S. Hydrogenotrophic culture enrichment reveals rumen Lachnospiraceae and Ruminococcaceae acetogens and hydrogen-responsive Bacteroidetes from pasture-fed cattle. FEMS Microbiol. Lett. 362, (2015).

30. Gagen, E. J. et al. Functional gene analysis suggests different acetogen populations in the bovine rumen and tammar wallaby forestomach. Appl. Environ. Microbiol. 76, 7785–7795 (2010).

31. Gagen, E. J. et al. Investigation of a new acetogen isolated from an enrichment of the tammar wallaby forestomach. BMC Microbiol. 14, 314 (2014).

32. Meyer, B. & Kuever, J. Molecular analysis of the diversity of sulfate-reducing and sulfur-oxidizing prokaryotes in the environment, using aprA as functional marker gene. Appl. Environ. Microbiol. 73, 7664–7679 (2007).

33. Devkota, S. et al. Dietary-fat-induced taurocholic acid promotes pathobiont expansion and colitis in Il10−/− mice. Nature 487, 104–108 (2012).

34. Denman, S. E., Tomkins, N. W. & McSweeney, C. S. Quantitation and diversity analysis of ruminal methanogenic populations in response to the antimethanogenic compound bromochloromethane. FEMS Microbiol. Ecol. 62, 313–322 (2007).

35. Stewart, R. D. et al. Compendium of 4,941 rumen metagenome-assembled genomes for rumen microbiome biology and enzyme discovery. Nat. Biotechnol. 37, 953–961 (2019).

36. Xie, F. et al. An integrated gene catalog and over 10,000 metagenome-assembled genomes from the gastrointestinal microbiome of ruminants. Microbiome 9, 137 (2021).

37. Carbonero, F., Benefiel, A. C. & Gaskins, H. R. Contributions of the microbial hydrogen economy to colonic homeostasis. Nat. Rev. Gastroenterol. Hepatol. 9, 504–518 (2012).

38. Gao, Z., Karlsson, I., Geisen, S., Kowalchuk, G. & Jousset, A. Protists: Puppet masters of the rhizosphere microbiome. Trends Plant Sci. 24, 165–176 (2019).

39. Caron, D. A., Worden, A. Z., Countway, P. D., Demir, E. & Heidelberg, K. B. Protists are microbes too: a perspective. ISME J. 3, 4 (2009).

40. Hook, S. E., Steele, M. A., Northwood, K. S., Wright, A. D. & McBride, B. W. Impact of high-concentrate feeding and low ruminal pH on methanogens and protozoa in the rumen of dairy cows. Microb. Ecol. 62, 94–105 (2011).

41. Towne, G., Nagaraja, T. G. & Kemp, K. K. Ruminal ciliated protozoa in bison. Appl. Environ. Microbiol. 54, 2733–2736 (1988).

42. Wallace, R. J. et al. A heritable subset of the core rumen microbiome dictates dairy cow productivity and emissions. Sci Adv 5, eaav8391 (2019).

43. Gast, R. J., Sanders, R. W. & Caron, D. A. Ecological strategies of protists and their symbiotic relationships with prokaryotic microbes. Trends Microbiol. 17, 563–569 (2009).

44. Pernthaler, J. Predation on prokaryotes in the water column and its ecological implications. Nat. Rev. Microbiol. 3, 537–546 (2005).

45. Paisie, T. K., Miller, T. E. & Mason, O. U. Effects of a ciliate protozoa predator on microbial communities in pitcher plant (Sarracenia purpurea) leaves. PLoS One 9, e113384 (2014).

46. Bernhard, J. M., Buck, K. R., Farmer, M. A. & Bowser, S. S. The Santa Barbara Basin is a symbiosis oasis. Nature 403, 77–80 (2000).

47. Ott, J., Bright, M. & Bulgheresi, S. Marine microbial thiotrophic ectosymbioses. Oceanogr. Mar. Biol. Annu. Rev. 42, 95–118 (2004).

48. Kuwahara, H., Yuki, M., Izawa, K., Ohkuma, M. & Hongoh, Y. Genome of ‘Ca. Desulfovibrio trichonymphae’, an H2-oxidizing bacterium in a tripartite symbiotic system within a protist cell in the termite gut. ISME J. 11, 766–776 (2017).

49. Ohkuma, M. et al. Acetogenesis from H_2_ plus CO_2_ and nitrogen fixation by an endosymbiotic spirochete of a termite-gut cellulolytic protist. Proc. Natl. Acad. Sci. U. S. A. 112, 10224–10230 (2015).

50. Ikeda-Ohtsubo, W. et al. ‘Candidatus Adiutrix intracellularis’, an endosymbiont of termite gut flagellates, is the first representative of a deep-branching clade of Deltaproteobacteria and a putative homoacetogen. Environ. Microbiol. 18, 2548–2564 (2016).

51. Graf, J. S. et al. Anaerobic endosymbiont generates energy for ciliate host by denitrification. Nature 591, 445–450 (2021).

52. Welty, C. M. et al. Rumen microbial responses to supplemental nitrate. II. Potential interactions with live yeast culture on the prokaryotic community and methanogenesis in continuous culture. J. Dairy Sci. 102, 2217–2231 (2019).

53. Roman-Garcia, Y. et al. Rumen microbial responses to supplemental nitrate. I. Yeast growth and protozoal chemotaxis in vitro as affected by nitrate and nitrite concentrations. J. Dairy Sci. 102, 2207–2216 (2019).

54. Williams, A. G. & Coleman, G. S. The rumen protozoa. (Springer Science & Business Media, 2012).

55. Belanche, A., de la Fuente, G. & Newbold, C. J. Study of methanogen communities associated with different rumen protozoal populations. FEMS Microbiol. Ecol. 90, 663–677 (2014).

56. Stevenson, D. M. & Weimer, P. J. Dominance of Prevotella and low abundance of classical ruminal bacterial species in the bovine rumen revealed by relative quantification real-time PCR. Appl. Microbiol. Biotechnol. 75, 165–174 (2007).

57. Caporaso, J. G. et al. Global patterns of 16S rRNA diversity at a depth of millions of sequences per sample. Proc. Natl. Acad. Sci. U. S. A., 4516–4522 (2011).

58. Tapio, I. et al. Oral Samples as Non-Invasive proxies for assessing the composition of the rumen microbial community. PLoS One 11, e0151220 (2016).

59. Yuan, J. S., Reed, A., Chen, F. & Stewart, C. N., Jr. Statistical analysis of real-time PCR data. BMC Bioinformatics 7, 85 (2006).

60. Katoh, K., Rozewicki, J. & Yamada, K. D. MAFFT online service: multiple sequence alignment, interactive sequence choice and visualization. Brief. Bioinform. 20, 1160–1166 (2019).

61. Fu, L., Niu, B., Zhu, Z., Wu, S. & Li, W. CD-HIT: accelerated for clustering the next-generation sequencing data. Bioinformatics 28, 3150–3152 (2012).

62. Minh, B. Q. et al. IQ-TREE 2: New models and efficient methods for phylogenetic inference in the genomic era. Mol. Biol. Evol. 37, 1530–1534 (2020).

63. Letunic, I. & Bork, P. Interactive Tree Of Life (iTOL) v4: recent updates and new developments. Nucleic Acids Res. 47, W256–W259 (2019).

64. Seemann, T. Prokka: rapid prokaryotic genome annotation. Bioinformatics 30, 2068–2069 (2014).

65. Aramaki, T. et al. KofamKOALA: KEGG Ortholog assignment based on profile HMM and adaptive score threshold. Bioinformatics 36, 2251–2252 (2020).

66. Kanehisa, M. & Goto, S. KEGG: kyoto encyclopedia of genes and genomes. Nucleic Acids Res. 28, 27–30 (2000).

67. Bolyen, E. et al. Reproducible, interactive, scalable and extensible microbiome data science using QIIME 2. Nat. Biotechnol. 37, 852–857 (2019).

68. Callahan, B. J. et al. DADA2: High-resolution sample inference from Illumina amplicon data. Nat. Methods 13, 581–583 (2016).

69. Quast, C. et al. The SILVA ribosomal RNA gene database project: improved data processing and web-based tools. Nucleic Acids Res. 41, D590–6 (2013).

70. Hammer, Ø., Harper, D. A. T. & Ryan, P. D. PAST: Paleontological Statistics Software Package for Education and Data Analysis. Palaeontol. Electronica 4, 9pp (2001).

71. van den Boogaart, K. G. & Tolosana-Delgado, R. ‘compositions’: A unified R package to analyze compositional data. Comput. Geosci. 34, 320–338 (2008).

72. Computing, R. & Others. R: A language and environment for statistical computing. R Core Team (2013).

