## Supplementary material for "Rumen protozoa are a hub for diverse hydrogenotrophic functions"

**Figure S1.** Abundance of prokaryotes and protozoa across animal diets.

**Figure S2.** Prokaryotic composition of the protozoa associated community.

**Figure S3.** Abundance of *Desulfovibrionaceae*.

**Table S1.** Primers used for community analysis, amplicon sequencing, and hydrogen utilization.
**Table S2.** ANOSIM analysis across the different prokaryotic communities and across the different diets.

**Table S3.** Genus abundances.

**Table S4.** Gene similarities.

### Supplementary figures

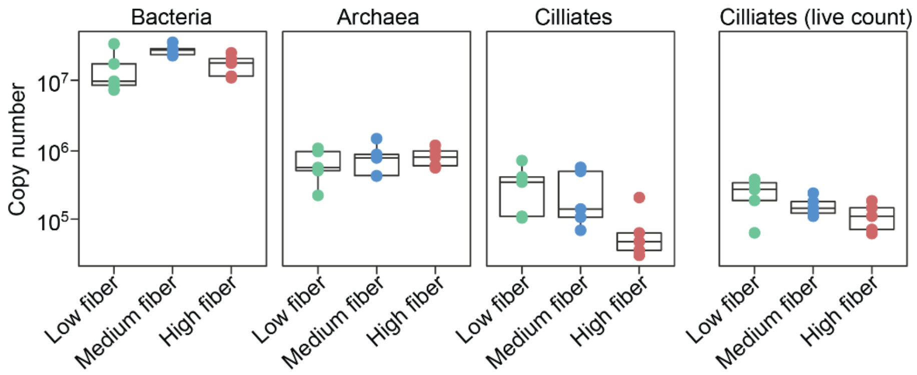


**Figure S1. Abundance of prokaryotes and protozoa across animal diets**. The abundance of bacteria, archaea and ciliate protozoa across different diets was quantified using quantitative PCR with the appropriate primers (see methods). The plot on the right shows the number of ciliate protozoa evaluated by counting protozoa cells under a microscope (see methods).

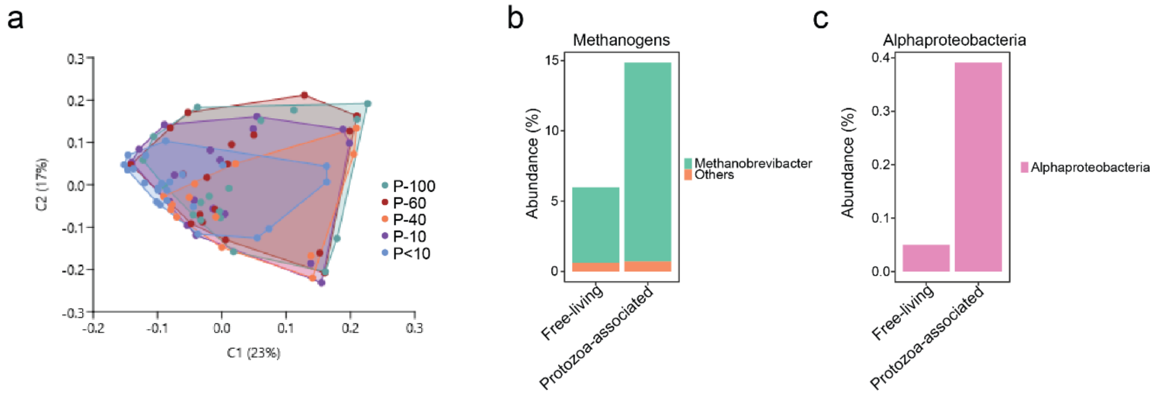


**Figure S2. Prokaryotic composition of the protozoa associated community.** (a) Principal coordinate analysis (PcoA) between different sub-communities of protozoa within each cow. (b), (c) Relative abundance of Methanogens and Alphaproteobacteria in the free-living community vs. the protozoa associated community.

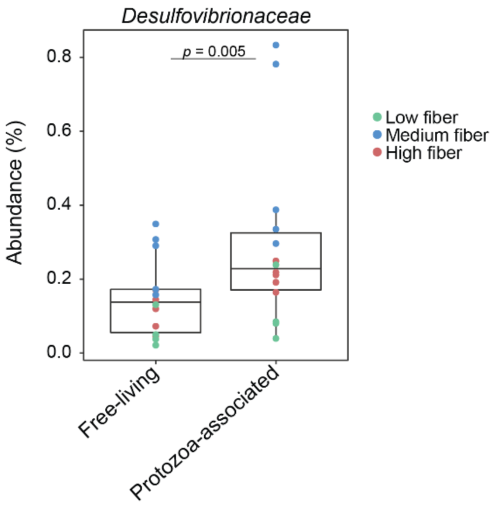


**Figure S3. Abundance of *Desulfovibrionaceae*.** The abundance of Desulfovibrionaceae was evaluated using 16s rRNA sequencing for free-living bacteria and protozoa-associated bacteria across the different diets. Paired Wilcoxon test was used to assess significance.

**Table S1.** Primers used for community analysis, amplicon sequencing, and hydrogen utilization

| Primer name | 5'-3' sequence | Size (bp) | Target | Reference |
|---|---|---|---|---|
| mcrA F | TTCGGTGGATCDCARAGRGC | 140 | Archaeal methyl coenzyme-M reductase | Denman et al. (2007) |
| mcrA R | GBARGTCGWAWCCGTAGAATCC | | | |
| apra-1 F | GCGCCAACYGGRCCRTA | 359 | *aprBA* operon of *Desulfovibrio Vulgaris* spp (sulfate reducing). | Meyer and Kuever (2007) |
| AprA-5 R | GCGCCAACYGGRCCRTA | | | |
| FTHFS F | TTYACWGGHGAYTTCCATGC | 1102 | Formyltetrahydrofolate Synthetase Gene (Acetogenic bacteria) | Gagen et al. (2010) |
| FTHFS R | GTATTGDGTYTTRGCCATACA | | | |
| NRFA F | ATGACCCGGATACCATG | 498 | NiR of *Selenomonas ruminatium* (Nitrate reducing) | Asanuma et al. (2015) |
| NRFA R | AGTGGGCTGGTCATCCA | | | |
| ACS F | CTYTGYCAGTCMTTYGCBCC | 416 | Acetogenic bacteria | Gagen et al. (2010) |
| ACS R | CCCATAAABCCYGGDGTYTG | | | |
| aprA F | TGGCAGATMATGATYMACGGG | 396 | Sulfate Reducing | Deplancke et al. (2000) |
| aprA R | GGGCCGTAACCGTCCTTGAA | | | |
| dsrA F | CCAACATGCACGGYTCCA | 162 | Sulfate Reducing | Devkota et al. (2012) |
| dsrA R | CGTCGAACTTGAACTTGAACTTGTAGG | | | |

**Table S2. ANOSIM analysis across the different prokaryotic communities and across the different diets.** The values on the lower left side of the table represent the R value denoting the difference between the different group (closer to 1 = more dissimilar) and the upper right side of the table the corresponding p-values.

| R/P | Free-living HF | Free-living LF | Free-living MF | Protozoa associated HF | Protozoa associated LF | Protozoa associated MF |
|---|---|---|---|---|---|---|
| Free-living HF | | 0.0015 | 0.843 | 0.0015 | 0.006 | 0.006 |
| Free-living LF | 0.7418 | | 0.12 | 0.0165 | 0.09 | 0.09 |
| Free-living MF | 0.2669 | 0.84 | | 0.006 | 0.132 | 0.09 |
| Protozoa associated HF | 0.8447 | 0.887 | 0.8831 | | 0.0345 | 0.1935 |
| Protozoa associated LF | 0.9535 | 0.824 | 0.932 | 0.5623 | | 0.04 |
| Protozoa associated MF | 0.9367 | 0.964 | 0.864 | 0.3478 | 0.592 | |

**Table S3. Genus abundances**. Relative abundance of prokaryotic genera in the free-living and protozoa associated community and under the different diets. Significance was obtained on the CLR transformed data using paired FDR corrected Wilcoxon test with the threshold value for significance at $p < 0.05$.

**Table S4. Gene similarities.** Similarity of clone sequences from this study to closest homolog in databases (Figure 4; see methods).
